# Coenzyme M: An Archaeal Antioxidant as an Agricultural Biostimulant

**DOI:** 10.1101/2024.12.20.629737

**Authors:** Jeremy H. Brown, Jithesh Vijayan, Aline Rodrigues de Queiroz, Natalia Figueroa Ramos, Nate Bickford, Melissa Wuellner, Nicole R. Buan, Julie M. Stone, Katarzyna Glowacka, Rebecca L. Roston

## Abstract

Rising global food demand necessitates improved crop yields. Biostimulants offer a potential solution to meet these demands. Among them, antioxidants have shown potential to improve yield, nutritional quality and resilience to climate change. However, large-scale production of many antioxidants is challenging. Here, we investigate Coenzyme M (CoM), a small, achiral antioxidant from archaea, as a potential biostimulant, investigating its effects on growth and physiology. CoM significantly increased biomass of the model plant *Arabidopsis thaliana* in a concentration-dependent manner consistent with it being metabolized. In tobacco, CoM increased photosynthetic light capture capacity, consistent with observed growth improvements. Interestingly, this effect was independent of carbon capture rates. Furthermore, CoM promoted early-stage shoot growth in various crops species, including tobacco, basil, cannabis, and soybean. Our results suggest CoM is a promising, scalable biostimulant with potential to modify photosynthesis and enhance crop productivity.

**Key Points:**

- Antioxidant application improves plant growth but is typically too expensive to use widely.
- Coenzyme M is a small, easily made antioxidant of Archaeal origin.
- Applied exogenously, Coenzyme M conditionally increases plant biomass.
- Coenzyme M represents an antioxidant with potential to be a widely used biostimulant.

## Introduction

Current models of population growth project a 35 to 56% increase in global food demand by 2050, necessitating improved crop yield efficiency to meet food security goals (van Dijk et al. 2021). Application of biostimulants directly to field crops or even specialty crops produced under controlled environments (greenhouse or hydroponics) may provide an ideal solution (Rouphael and Colla 2020). One mode of action of biostimulants is through the modulation of reactive oxygen species (ROS) (Hasanuzzaman et al. 2021). As such, antioxidants can act as biostimulants, and their capacity to confer conditional growth benefits have been reported across a wide variety of plant species under an array of environmental stress conditions (Jan et al. 2022, Khalid et al. 2022, Rodrigues de Queiroz et al. 2023). Mechanisms for how antioxidants increase plant growth remain unclear, however, modification of redox homeostasis and amelioration of oxidative stress are hypothesized to be primary determinants in facilitating the growth enhancements attributed to exogenous antioxidants (Khalid et al. 2022, Rodrigues de Queiroz et al. 2023). Additional rationales for improved growth suggest that applied antioxidants increase photosynthetic output by interacting with redox-responsive photosynthetic components. Indeed, modulation of photosynthetic parameters has been observed in response to exogenous application of antioxidants under control and stress conditions (Rodrigues de Queiroz et al. 2023, Zhou et al. 2018).

Photosynthesis has long been a target for improved crop efficiency, as improvements to photosynthetic capacity are often linked to improved productivity and yield (Kromdijk et al. 2016, López-Calcagno et al. 2020). Photosystem II (PSII) is often the target for photosynthesis improvements since it is responsible for primary light capture. The maximum quantum yield (*F*_*v*_/*F*_*m*_) and efficiency of PSII (□*PSII*) are parameters that are affected strongly by the buildup of ROS, which can occur readily under abiotic stresses such as high-light, where PSII is unable to accommodate for excess electrons from light capture (Pospíšil 2016). Nonphotochemical quenching (NPQ) of chlorophyll fluorescence is one way that plants have to disperse excess energy in the form of heat prior to the buildup of ROS, although the specific mechanisms for NPQ are not fully elucidated (Ruban and Wilson 2021). Excess energy not dissipated by NPQ or other means results in the generation of ROS within the chloroplast, which must be quenched or will result in the disassembly of light harvesting complexes through oxidative damage (Fristedt and Vener 2011). This ROS can be quenched through antioxidant pathways like the ascorbate-glutathione cycle (Li and Kim 2022). The results of PSII energy use is the production of a proton gradient, ATP, and reducing equivalents, which are used in the Calvin-Bensen-Bassam cycle to fix carbon dioxide and transform captured light energy into chemical energy (Croce et al. 2024). Application of antioxidants has caused variable effects on photosynthetic fluorescent parameters, but there is less information on how antioxidant application affects gas exchange and carbon fixation (Ding et al. 2016, Huang et al. 2019).

Coenzyme M (CoM) is a small antioxidant isolated from archaea, where it serves as a key component for methane production via the Wolfe cycle (Thauer 2012). CoM’s structure is comprised of a sulfonate group connected to a thiol by a two-carbon bridge (Fig. 1A) and is widely used as a human cancer medication under the drug name Mesna (Links and Lewis 1999). Mesna/CoM serves as a chemoprotectant against acrolein, a urotoxic metabolite that can be excreted into the urinary tract in response to some chemotherapy treatments. Mesna/CoM’s potent chemoprotective property stems from CoM’s reactivity with alkyl groups, forming inactive thioethers in the process. CoM is achiral, rendering it relatively easy to produce in abundance through chemical synthesis with 1,2-dichloroethane as the starting material (Dao and Nguyen 2018). Its structural simplicity and synthetic scalability position CoM as a candidate for agricultural applications, particularly when evaluated against natural antioxidants like glutathione (GSH), which share similar properties but face production limitations.

**Figure 1.**
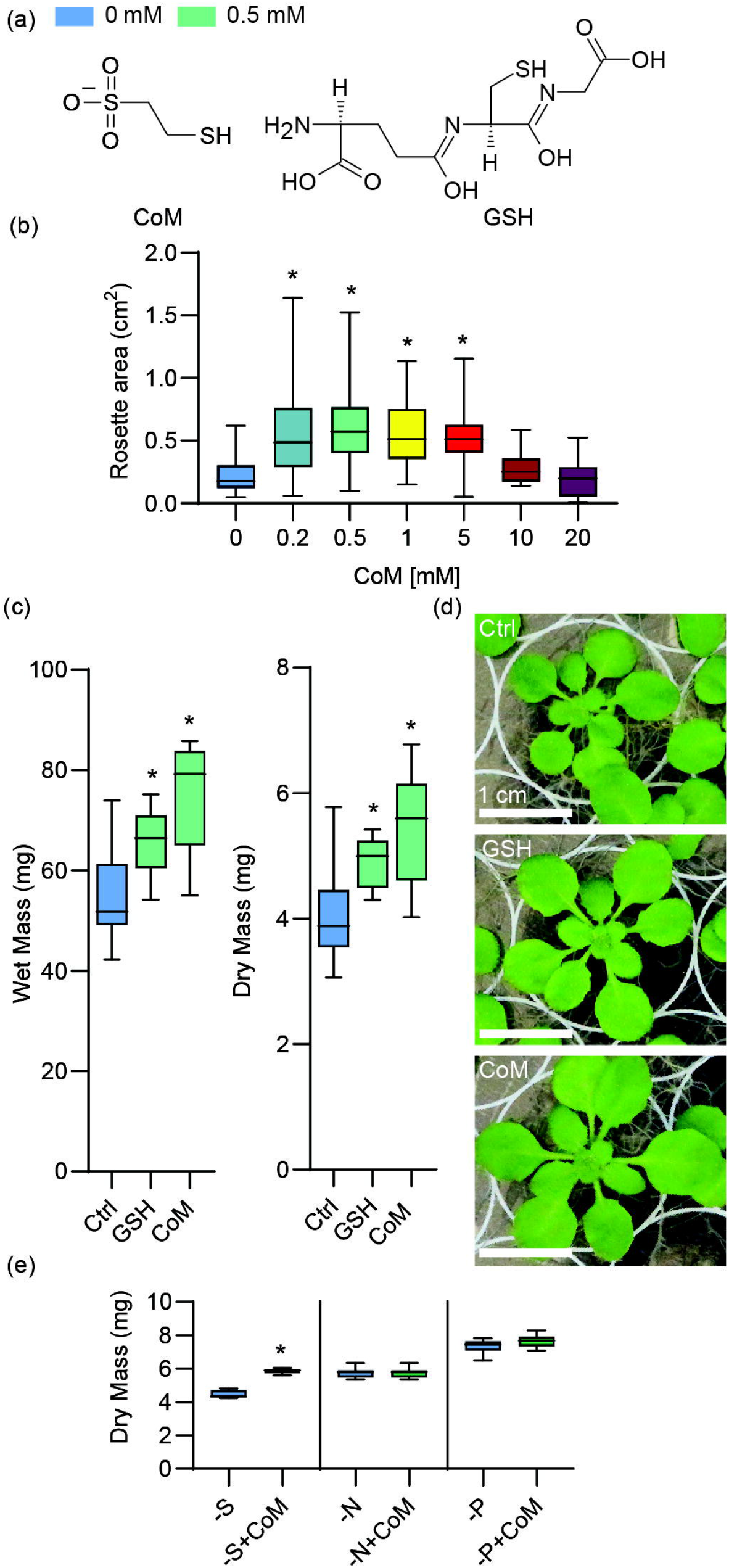
Growth and nutrient limitation effects of Coenzyme M application in Arabidopsis. (a) Structure of Coenzyme M (CoM) and Glutathione (GSH). (b) Rosette area measurements of Arabidopsis at 25 days; grown on full-strength MS supplemented with CoM, n ≥ 41 biological replicates over ≥ 2 trials (c) Dry mass measurements of Arabidopsis grown for 21 days on solid full-strength or half-strength MS media supplemented with 0.5 mM CoM or GSH. Mass on a per plant basis with four measurements per trial for three trials, n ≥ 8 biological replicates. (d) Photographs of Arabidopsis from (c), scale bar is 1 cm. (e) Dry mass of Arabidopsis grown for 21 days with or without 0.5 mM CoM on half-strength MS media formulated without Sulfur, Nitrogen, or Phosphorus as indicated. n ≥ 7 biological replicates. Asterisks indicate p-value ≤ 0.05.

CoM is of archaeal origin, and not endogenously produced in humans nor plants. The closest comparable endogenous antioxidant to CoM in plants is Glutathione (GSH), which also contains a thiol group to confer redox activity (Fig. 1a). CoM and GSH share similar reduction potentials of -271 and -264 mV, respectively (Laird et al. 2024, Rost and Rapaport 1964). However, GSH is a chiral tripeptide, making it difficult to produce in abundance (Huang and Yin 2020). A major hurdle in the widespread use of antioxidants in agriculture is that large-scale antioxidant(s) synthesis/production can be challenging, due to the fact that effective antioxidants often possess multiple chiral centers hindering mass production (Hugel et al. 1995, Huang and Yin 2019, Reichstein and Grüssner 1934). If CoM has similar positive effects on growth of various plant species as has been reported for GSH (Rodrigues de Queiroz et al. 2023), it could be an economical, readily synthesized, exogenously applied effective alternative to increase plant growth, resilience, and crop yield. In this study we investigate the effects of CoM application on a multitude of plant species to determine its potential as an agricultural biostimulant. CoM was applied to the model species *Arabidopsis thaliana* as well as multiple agriculturally relevant species. Improved plant growth, including aerial and root tissue biomass and plant height were observed in response to exogenous CoM application. Moreover, CoM treatment elicited significant changes in a range of photosynthetic parameters that may contribute to our observed growth enhancements.

## Materials and Methods

### Plant growth

Arabidopsis: *Arabidopsis thaliana* (Columbia ecotype) was grown on solid full-strength or half-strength Murashige and Skoog (MS) with vitamins (Caisson Labs) media (pH 5.7) supplemented with CoM (AstaTech) or GSH (Acros Organics) at various concentrations. For nutrient limitation experiments, formulations of MS without N, P, or S (Caisson Labs) were used. Seeds were bleach sterilized, rinsed, and sown on media plates. Plates were placed in 4°C for 48 hours to break seed dormancy and then transferred to a growth chamber with a 16 h light / 8 h dark photoperiod with 22°C day at 150 μmol m^-2^ s^-1^ light, 18°C night.

Tobacco: *Nicotiana tabacum* was grown on soil under greenhouse conditions. Seeds were sown on a greenhouse mixed soil (8:8:3:1 (w/w/w/w) peat moss:vermiculite:sand:screened topsoil, with 7.5:1:1:1 (w/w/w/w) Waukesha fine lime, Micromax, Aquagro, and Green Guard per 0.764 m^2^). Plants were grown in a greenhouse with a 16 h light / 8 h dark photoperiod with 23-25°C day and 18-20°C night.

Basil: *Ocimum basilcum* v. Genovese (basil) seeds were germinated on wet paper towels and transplanted to greenhouse mixed soil after radicle emergence within two days. Plants were grown in a greenhouse with a 16 h light / 8 h dark photoperiod with 20-23°C day and 18-20°C night.

Cannabis: *Cannabis sativa* plants were clonally propagated from a single individual for the study and grown in greenhouse mixed soil. Greenhouse supplemental lighting was provided to generate a 16 h light / 8 h dark photoperiod with temperatures set between 20-23°C in the day and 18-20°C in the night.

Soybean: *Glycine max* (Thorne) seeds were sown on a greenhouse mixed soil. Plants were grown in a greenhouse with a 16 h light / 8 h dark photoperiod with 23-25°C day and 18-20°C night. Plants were grown to 21 days after sowing with biweekly spray treatment beginning at 7 days post sowing.

### CoM Application

Solid media: In experiments where plants were grown on solid MS media, filter-sterilized (0.2 μ m) CoM or GSH dissolved in water were added to autoclaved MS media solutions immediately prior to solidification.

Spray: In experiments where CoM was applied via spray application, fresh solutions of CoM in MilliQ water were prepared immediately prior to application. CoM solutions were sprayed to cover the majority of the adaxial surfaces of the leaves and allowed to dry in the plant’s growth chambers.

Seed soak: In experiments where CoM was applied via seed soaking, solutions of CoM in MilliQ water at concentrations indicated in figure legends were prepared and seeds were soaked at room temperature overnight (14 hours) prior to planting.

### Plant Tissue and Size Measurements

Tissue Collection: Aerial shoot and leaf tissues were collected and weighed to measure fresh mass, then tissue was dried at 80°C for 3-4 days to measure dry mass. For Arabidopsis experiments on solid media, multiple individual plants were aggregated, and data is presented on a per-plant basis. Measurements for tobacco, soybean, and other species are represented on the basis of individual plants.

Rosette size measurements: Top-down photos of Arabidopsis were taken with a 1 cm^2^ size marker. Rosette size was calculated using Easy Leaf Area (Ealson and Bloom 2014).

Overlapping plants were removed at 25 days of growth.

Height and area measurements: Height from the base of the stem to the highest point of the aerial tissues was recorded. To determine approximate area, plant height was multiplied by the width of aerial tissues at their widest point.

### Sulfur analysis

Arabidopsis shoot tissue was collected after 21 days of growth and 50 mg of aerial tissue was used for ICPMS analysis. Intracellular sulfur content measured by inductively coupled plasma mass spectrometry (ICPMS) as described in Seravalli 2012 and Seravalli 2024.

### Photosynthetic analyses

To measure chlorophyll fluorescence, plants were submitted to a pulse-amplitude modulated protocol during 10 min of light illumination at 800□μmol□m^−2_^s^−1^ followed by 10□min of darkness using a chlorophyll fluorescent imager (FluorCam FC 800-C). Saturating flashes of 2,400□μmol□m^−2^□s^−1^ for a duration of 800□ms were used in the light and dark periods. First, *F*_*o*_ was measured in dark-adapted plants, followed by capture changes in steady-state fluorescence (*F*_*s*_) and maximum fluorescence under illuminated conditions (*F*_*m*_′) over time, the measurements were done at the following time points 0, 0.25, 0.5, 0.75, 1, 2, 3, 4, 5, 6, 7, 8, 9, 10, 10.25, 10.5, 11, 12, and 15 min. The raw data were processed automatically with fluorescence background exclusion. Maximum quantum yield is defined as *F*_*v*_/*F*_*m*_ in dark-adapted plants, and NPQ is defined as *F*_*m*_/(*F*_*m*_*’*-1). The data was fitted to hyperbola equations according to Sahay et al. (2023) where the induction rate Ind_(Rate)_ was the initial slope of the hyperbola in light, and the relaxation rate Rel_(Rate)_ was the initial slope hyperbola in the dark. For our dose-response curves, we performed a baseline correction to determine the percent change in those parameters relative to control measurements. Those results were fitted into a polynomial equation.

Gas exchange measurements were collected according to Sahay et al. (2023) with the exceptions that block temperature was set to 25°C, [CO_2_] was set to 400 μmol, water vapor pressure deficit was set to 1.3kPa, and maximum light intensity was set to 2000 μmol m^-2^ s^-1^.

### Statistical analyses

All statistical analyses were performed using Graphpad Prism. As indicated in Table S1, ANOVAs and corrected t-tests were performed as appropriate. To compare pair-wise values, one-way ANOVAs were followed by Dunnett’s multiple comparisons tests, or two-way ANOVAs were followed by Bonferroni’s multiple comparison test. Finally, for experiments with only two comparison groups, Welch’s corrected t-test, 2-tailed, and unpaired was used. Throughout the paper, p-values represent the adjusted p-value determined by the indicated statistical test with the number of biological replicates represented (n).

## Results

### Coenzyme M improves growth and is metabolized by Arabidopsis

As an initial investigation into CoM’s potential applicability to improve plant biomass, we used the model plant *Arabidopsis thaliana* ecotype Columbia (Arabidopsis). To determine optimal concentrations of CoM for application, seeds were grown on plates of solid synthetic growth media supplemented with CoM concentrations from 0.2 to 20 mM (Fig. 1b). Results showed that concentrations of CoM between 0.2 and 5 mM significantly increased the rosette size of Arabidopsis at 25 days of growth (Fig. 1b). The maximal enhancement reached was 168% of untreated control plants with 0.5 mM CoM at 25 days (p-values in Table S1). The presence of CoM did not appear to affect germination (Fig. S1).

Similar to CoM, another thiol-based reductant, glutathione (GSH), has been applied to plants at comparable concentrations (0.05 to 1 mM) to investigate its impact on growth (Nakamura et al. 2020, Kim et al. 2017, Cheng et al. 2015). To directly compare the effects of CoM and GSH, we assessed Arabidopsis biomass after growth in two common synthetic media, full-strength and half-strength Murashige-Skoog (MS, Pasternak et al. 2023), supplemented with 0.5 mM of either reductant. Both reductant treatments caused increased mass of aerial tissues at 21 days of growth relative to mock treatment (Fig. 1c, d). The dry mass increased by 35% and 22% with CoM and GSH supplementation, respectively, in full MS medium. Fresh mass was increased similarly. Under half-strength MS conditions, the increase in Arabidopsis aerial tissue was only significant in fresh mass, 30% for CoM and 17% for GSH, suggesting a media-dependent effect. To assess the root growth response to CoM, Arabidopsis were grown on vertical plates supplemented with 0.5 mM CoM or GSH. Again, both antioxidants increased root length versus control, by 47% or 34% for CoM or GSH respectively (Fig. S2).

GSH is a naturally occurring antioxidant in plants, while CoM is not, but they both had similar impacts on growth in our assays. We wanted to understand the likelihood that CoM, like GSH, is taken up and metabolized (Arianmehr et al 2022). Because CoM contains sulfur, we investigated the possibility that it could provide this nutrient, by testing the growth of plants in CoM supplementation in MS media reformulated to lack each of the major macronutrients, nitrogen, phosphorus, and sulfur. Arabidopsis grown on nitrogen- and phosphorus-limited media showed no significant changes with CoM supplementation, while those grown on sulfur-limited medium showed an increase in dry mass of 30.2% when supplemented with 0.5 mM CoM (Fig. 1e), suggesting that CoM was taken up and metabolized for sulfur. However, direct measurement of sulfur content in plants supplemented with CoM or GSH revealed no detectable differences compared to untreated controls (Fig. S3), suggesting that supplementation supports growth under sulfur limitation without measurable increases in sulfur at this developmental stage.

### Coenzyme M increases photosynthetic capacity

Increased plant growth is often linked to changes in photosynthetic parameters (Kromdijk et al., 2016; Hubbart et al., 2018; De Souza et al., 2022; Xin et al., 2023), therefore, to investigate whether CoM treatment influenced photosynthesis, we conducted greenhouse trials on *Nicotiana tabacum* (tobacco), a common model for photosynthesis studies (Ramzan et al.2021, Ding et al. 2016, Huang et al. 2019). We began by verifying the effect of CoM on growth.

Tobacco sprayed with CoM at concentrations between 0.5 to 5 mM significantly increased biomass in a concentration-dependent manner (p = 0.0114, one-way ANOVA). (Fig. 2a, Fig. S4. The maximum increase of arial tissue dry mass was 48% with 1 mM spray application, with a second trial confirming the trend of 1 mM resulting in the most significant increase in plant biomass (Fig. S4). To determine whether CoM treatment influenced photosynthesis, we examined multiple parameters in dark-adapted tobacco sprayed with CoM at concentrations of 1, 2.5, and 5 mM. The maximum efficiency of photosystem II (*F*_*v*_/*F*_*m*_) increased with CoM treatment by 7.6% with 1 mM CoM (p = 0.0001, one-way ANOVA; Fig. 2b). *F*_*v*_/*F*_*m*_ is calculated from multiple components, thus we checked if minimum (*F*_*0*_), maximum (*F*_*m*_), or variable fluorescence (*F*_*v*_, *F*_*p*_) were the largest contributing component. We found that *F*_*m*_, *F*_*v*_ and *F*_*p*_ each increased after CoM treatment, consistent with an increase in the number of open photosystem II reaction centers (Fig. 2c). We then challenged dark adapted plants with an actinic light and pulses to saturate photosynthetic capacity and activate regulated energy dissipation measured as non-photochemical quenching (NPQ) followed by dark relaxation. Under these conditions, application of CoM increased the operating quantum yield of photosystem II (□*PSII*) at the end of both the light (Fig. 2d2) and dark (Fig. 2d4), though it did not affect the rates of induction (Fig. 2d1) or relaxation (Fig. 2d3). Similarly, CoM increased the maximum NPQ in the light (Fig 2e2), the rate of NPQ relaxation (Fig. 2e3) and residual NPQ in the dark (Fig 2e4). The observed increases of open photosystem II reaction centers (*F*_*m*_, *F*_*v*_ and *F*_*p*_), yield of photosystem II (□*PSII*) and increased rate of NPQ relaxation are all consistent with increased energy capture.

**Figure 2.**
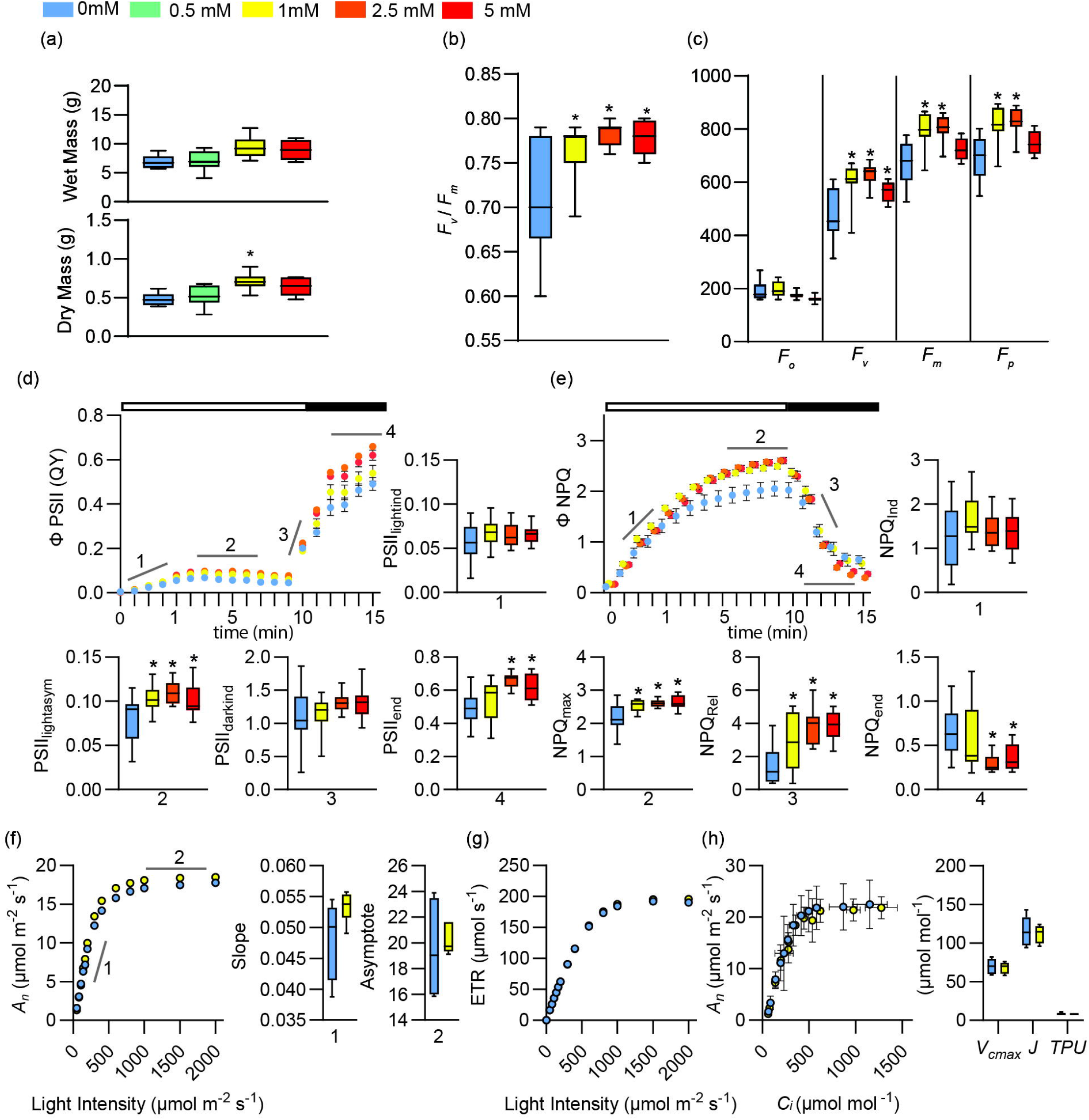
Photosynthetic parameter effects of Coenzyme M application in tobacco. (a) Fresh and dry mass measurements of tobacco grown on soil for 21 days with aerial tissues sprayed with CoM twice weekly, n = 6 biological replicates. Identically grown tobacco plants were tested for their photosynthetic traits as measured by FluorCam FC 800-C in panels (b – h). (b) The maximum quantum yield, *F*_*v*_/*F*_*m*_, measured after dark adaptation, n = 12 biological replicates. (c) Measurements of minimum (*F*_*0*_), maximum (*F*_*m*_), or variable fluorescence (*F*_*v*_, *F*_*p*_), after dark adaptation, n =12 biological replicates. (d) Photosystem II operating efficiency (□PSII) measured after biweekly spray of water, 1mM, 2.5mM, or 5mM CoM, and dark adaptation. Component traits of □PSII are indicated by grey lines and numbers and represented in separate graphs: (1) PSII_lightind_, the initial slope of a linear fit including the first two points and zero in the light (indicated by the white bar at the top of the □PSII graph), (2) PSII_lightasym_, the hyperbolic asymptote during the light, (3) PSII_darkind_, the slope of the first two points of PSII induction in the dark (indicated by the black bar at the top of the □PSII graph), and (4) PSII_end_, the hyperbolic asymptote during the dark, n = 12 biological replicates (e) Nonphotochemical quenching (□NPQ) measured under the same conditions as □PSII, and its component traits indicated by grey lines and numbers and represented in separate graphs, (1) NPQ_ind_, the initial slope of a linear fit including the first two points and zero in the light, (2) NPQ_max_, the hyperbolic asymptote during the light, (3) NPQ_rel_, the slope of the last light point and first two points of NPQ relaxation in the dark, and (4) NPQ_end_, the hyperbolic asymptote during the dark, n = 12 biological replicates. Panels (f – g) represent LiCOR 6800 measurements of 21-day-old tobacco plants sprayed biweekly with 1mM CoM. (f) Carbon assimilation (*A*_*n*_) in response to increasing light, and its component parts, the initial slope (1) and hyperbolic asymptote (2), n = 6 biological replicates. (g) The electron transport rate (ETR), n = 6 biological replicates. (h) The internal concentration of CO_2_ (*C*_*i*_) in response to increasing light, and maximum carboxylation rate allowed by Rubisco (*V*_*cmax*_), the rate of photosynthetic electron transport based on NADPH requirement (*J*), and triose phosphate use (*TPU*) fit from the *C*_*i*_ curve. All asterisks indicate p-value ≤ 0.05.

To explore the effect of CoM on carbon capture, we measured gas exchange parameters in response to increasing light and CO_2_ concentrations using an open gas exchange system (LI-COR 6800). We found that tobacco treated with CoM seemed to have a slight increase relative to controls in net carbon assimilation (*A*_*n*_) during both light-limited (slope) and light-saturated (asymptote) portion of the curve, though the effect was too small to be statistically significant at the 95% confidence interval (Fig. 2f). We also checked for CoM treatment impacts to the electron transport rate or CO_2_ usage and found neither were affected (Fig. 2h). Specifically, we measured how carbon assimilation varied with the change of CO_2_ dosage, and estimated limiting factors by curve fitting (Sharkey et al. 2007), defining the effect to maximum carboxylation rate allowed by Rubisco (*V*_*cmax*_), the rate of photosynthetic electron transport (*J*), and triose phosphate use (*TPU*), none of which were impacted by CoM treatment. Stomatal conductance (*G*_*s*_) was not significantly affected by CoM application (Fig. S5).

### CoM improves growth of a range of crop species

Model plants Arabidopsis and tobacco both showed improved growth as a result of CoM application, therefore we applied CoM to more agriculturally relevant species to explore potential use in an agricultural capacity. In two trials we used *Ocimum basilcum* v. Genovese (basil, Fig. 3a, b). Basil showed a height increase after 28 (98%) and 35 days (111%) of growth when seeds were soaked with 0.5 mM CoM and sprayed once weekly with 0.5 mM CoM (Fig. 3a). To test the concentration-dependence of CoM, CoM was applied to basil by foliar spray twice weekly with concentrations between 1 to 5 mM. The greatest increase in basil dry mass was observed after treatment with 2 mM CoM (40%; Fig. 3b). *Cannabis sativa* (cannabis) received a foliar spray with 3 mM CoM three times weekly. Height was increased at 28 days (25%) and 35 days (38%) of growth relative to control plants (Fig. 3c). The shoot area of these plants was also increased at 35 days of growth (69%, Fig. 3d). To broaden the potential applicability of CoM treatment to commodity crops we also tested *Glycine max* v. Thorne (soybean). After leaflets emerged, they received a foliar spray of 0.5 or 1 mM CoM twice a week. Aerial tissue dry mass was increased (17%) after 21 days of growth (stage V2 - V3) with 1 mM CoM application (Fig. 3e). We concluded that CoM application stimulated growth of multiple species following frequent foliar spray.

**Figure 3.**
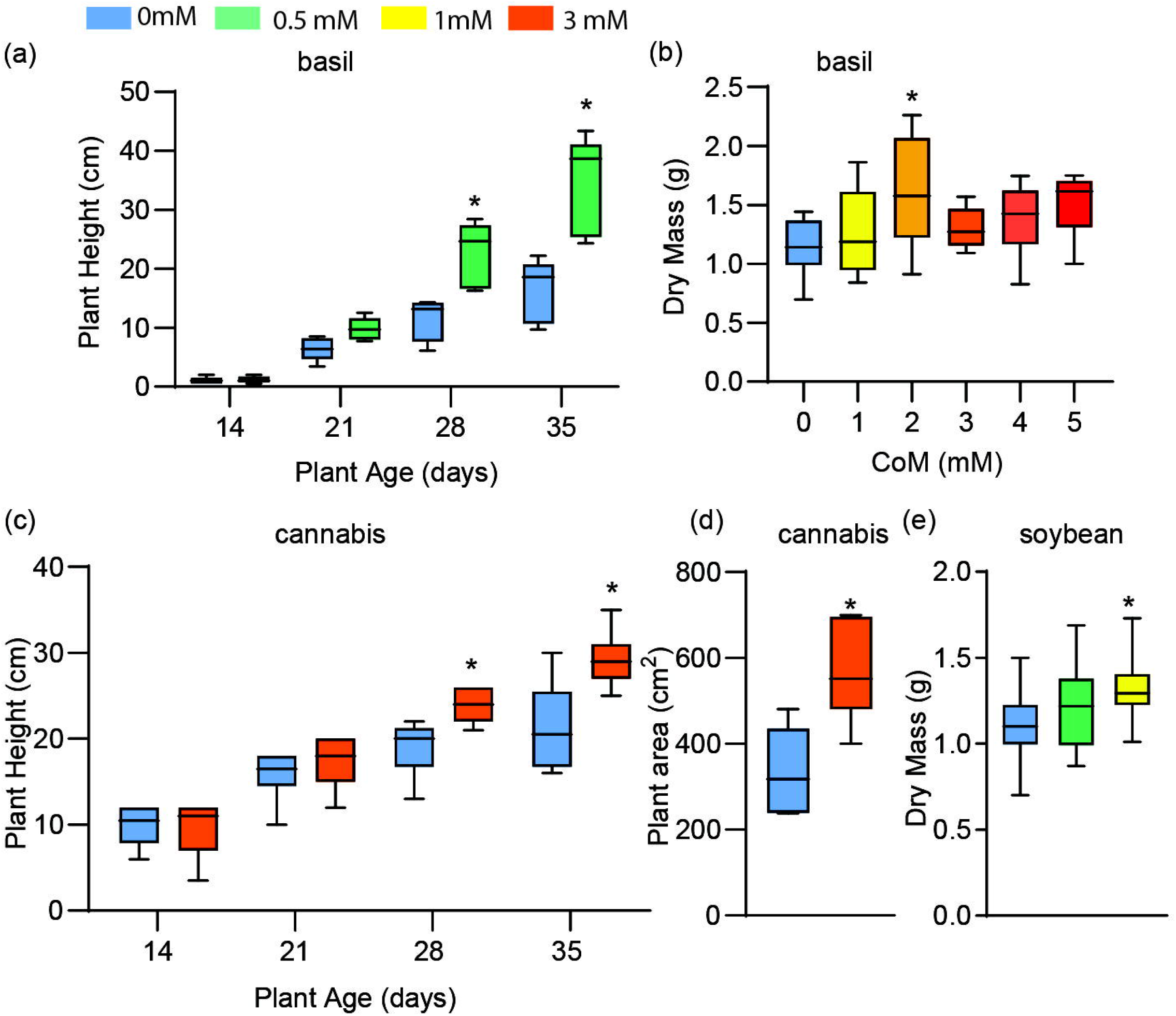
Growth effects of Coenzyme M application in various plant species - basil, cannabis, and soybean. (a) Plant height measurements of basil at ages indicated below. Seeds were soaked in 0.5 mM CoM prior to sowing and plants were sprayed once weekly with 0.5 mM CoM, n = 5 biological replicates. (b) Dry mass measurements of basil after 30 days of growth during which CoM at concentrations indicated below was sprayed three times a week, n = 9 biological replicates (c) Plant height measurements of cannabis at indicated ages that were sprayed three times weekly with 3 mM CoM, n = 6 biological replicates (d). Plant area measurements of cannabis sprayed three times weekly with 3 mM CoM, n = 6 biological replicates. (e) Dry mass measurements of soybean seedlings after 21 days of growth on soil during which aerial tissues were sprayed with CoM twice weekly, n = 30 biological replicates. All asterisks indicate p-value ≤ 0.05 All mass measurements represent aerial tissues.

## Discussion

Our results showed that the application of CoM enhanced growth in a species-specific and concentration-dependent manner. Arabidopsis, tobacco, basil, cannabis, and soybean all showed increased growth parameters with at least one concentration of CoM tested. Additionally, multiple growth parameters were affected by CoM application including wet and dry mass of aerial tissues (Fig 1c, 2a, 3b, and 3e), shoot height (Fig 3a and 3c), leaf area (Fig 1b and 3d), and root growth (Fig. S2). Many of these parameters were increased similarly in diverse plant species. In tobacco, we determined that CoM was directly influencing light energy capture, a likely cause of improved growth (Fig 2b, 2c, 2d, and 2e). Using CoM to increase the growth of aerial leaf tissues would be particularly beneficial for crop species like tobacco and basil, since their yield is leaf tissue. It is also likely to benefit soybean and other seed crops in which final yield is related to initial growth (Hilty et al. 2021).

### Mode of CoM action

Antioxidants may have different roles on the surface of plants and after internalization (Rodrigues de Queiroz et al. 2023). Since the ability of plants to internalize CoM was unknown, we investigated its potential to provide a source of sulfur, finding that CoM improved biomass of Arabidopsis grown in media lacking S, but not media lacking N or P (Fig, 1e). This is consistent with CoM being internalized and metabolized as a source of sulfur, though we cannot rule out the possibility that this effect is indirect. We attempted to measure the internal concentration of sulfur, however neither GSH nor CoM treatment measurably increased the sulfur content of Arabidopsis in our growth conditions (Fig. S3). GSH is generally considered to be internalized by Arabidopsis, due to the increase of internal GSH after application (Cheng et al 2015) and the phenotypic similarity of external application and internal upregulation (Chen et al. 2012). Since GSH application did not result in measurable S increases, and previous studies have found that GSH application does not always increase measurable sulfur depending on species, we conclude that CoM’s role as a metabolizable sulfur source remains plausible (Arianmehr et al. 2022).

Since photosynthesis is strongly affected by redox status and previous studies have found that antioxidant application can influence photosynthetic parameters, we investigated how CoM affects photosynthesis as a mode of action for increased growth (Ding et al. 2016, Huang et al. 2019). Application of CoM increased the maximum yield of photosystem II in both light and dark conditions (Fig. 2b, 2d) (Fig. 2c) but interestingly did not influence carbon assimilation in a significant way (Fig. 2f). These effects are consistent with the increase in tobacco growth (Fig. 2a). Additionally, NPQ_max_ and NPQ relaxation rate were improved with CoM application (Fig. 2e), suggesting a greater capacity to dissipate excess energy from light capture and return to a more efficient energy utilization state faster. Effects on NPQ rate were previously linked to growth increases (Kromdijk et al 2016). It is also an interesting result in the context of antioxidant application, as previous studies have shown that antioxidants have variable effects on NPQ (Ding et al. 2017, Rodrigues de Queiroz et al. 2023, Trojak and Skowron 2021). CoM application on tobacco simultaneously increased the plant’s ability to perform photosynthesis by increasing the amount of active PSII centers while also increasing their capacity to dissipate excess energy associated with high light in the form of NPQ, an uncommonly reported dual benefit for antioxidant application. Unexpectedly, these changes are independent of rubisco carboxylation (*V*_*cmax*_) and electron transport rate (*J*) (Fig. 2g, 2h), which other antioxidants have influenced (Rehman et al. 2021, Son et al. 2014). We conclude that changes in photosynthesis caused by application of CoM are consistent with observed increases to growth parameters and that CoM affected photosynthesis uniquely compared to application of other antioxidants like GSH (Ding et al. 2016).

### CoM impacts on growth

Benefits derived from the application of antioxidants are almost always concentration dependent (Zhang et al. 2015, Rodrigues de Queiroz et al. 2023). Applying a substance, antioxidant or otherwise, in sufficient concentration will have detrimental effects on growth and health of the plant treated (Kumari et el. 2022). For instance, melatonin applied at 100 mM inhibited root growth of *Brassica juncea*, but when it was applied at 0.1 mM it promoted root growth (Chen et al. 2009). We found CoM in concentrations between 0.2 and 5 mM had a positive growth benefit in Arabidopsis while concentrations of 10 mM and above had neutral or detrimental effects (Fig 1d). Effective concentrations appear to be species specific, as tobacco and soybean did not share an effective concentration of CoM (Figs 2a, 3e). Interestingly, when looking at agriculturally relevant species, CoM had a narrower range of effective concentration (Figs 2a, 3b, and 3e) compared to Arabidopsis (Fig 1b). This could be a byproduct of application method, as we applied CoM to Arabidopsis through media supplementation, while other species received a foliar spray.

### Potential for CoM use as a biostimulant

Agriculturally, biostimulants have been applied to seeds, plants, and soil through seed soaking or coating, foliar spray, or soil dry or wet applications, with the most common of these being foliar spray (Johnson et al. 2023). We mainly explored CoM application by foliar spray to agriculturally relevant species (Fig 2, 3). We found that it could provide a growth increase in tobacco up to 48%, in basil up to 39.9%, and in soybean up to 17.3% (Figs 2a, 3b, 3e). It’s important to note that the effective concentration of a biostimulant will be influenced by which plant it is being applied to and the method by which it is being applied (Ruzzi et al. 2024, Rodrigues de Queiroz et al. 2023). For CoM, we found that the most effective concentrations differed between tobacco and basil (Figs 2a and 3b). Previous studies with biostimulants also showed that environmental conditions can have a drastic effect on the efficacy of biostimulant application (Li et al. 2022, Sleighter et al. 2023) Trial-to-trial variation in tobacco suggested that CoM affects an aspect of plant growth that fluctuated with its environment. For example, two growth trials separated by three months resulted in different increases in dry mass, with one at 48% (Fig. 2a) and the other at 15.1% (Fig. S4). These trials also had different dry masses of control plant growth, presumably due to seasonal differences in day length and temperature, which is known to impact growth in the presence of supplemental lighting and heating (Matsubara 2018, Solbach et al. 2021).

ROS and the maintenance of ROS homeostasis are key signaling components for many growth processes (Mittler et al 2022). As such, altering ROS homeostasis through the addition of an antioxidant could introduce problems through the disruption of ROS signaling. An important consideration for the viability of an antioxidant as a biostimulant is whether it disrupts germination, since germination is heavily influenced by redox signaling (Bailey 2019). We found that CoM did not detrimentally affect the germination of Arabidopsis sown on media containing 0.5 mM CoM.

Finally, ease of synthesis is an aspect that is important to the economic value and availability of a biostimulant (Xu and Geelan 2018). GSH is a chiral molecule, and synthesis can be achieved through an enzymatic method in yeast which allows for the purification of GSH with the correct chirality (Huang and Yin 2019). Since CoM does not have chiral centers, it can be chemically synthesized without a biological host, and thus at larger scales. One method of synthesizing CoM uses 1,2-dichloroethane as starting material, which can readily be synthesized from ethanol (Dao and Nguyen 2018).

The ability to produce CoM in large quantities, lack of disruption in germination and health of plants, and ability to improve growth suggests that CoM has potential to be an effective and low cost biostimulant.

## Author Contribution Statement

JB, JV, ARQ, KG, NB, JMS, NRB, and RLR designed the research. NFR, JB, JV, ARQ, and NB performed the research and analyzed data. JB wrote the paper, all authors edited the paper.

## Acknowledgements

The authors gratefully acknowledge Samantha Link and Kandy Hanthorn for greenhouse services, Javier Seravalli for ICPMS analysis, and Ngoc Pham, Hailey Olberding, and Tom Ramaekers for their technical assistance and contributions to related research projects.

