## Supplementary material for "Coenzyme M: An Archaeal Antioxidant as an Agricultural Biostimulant": Table S1

**Figure S1.** **Germination rate of Arabidopsis is unaffected by CoM application** Germination rates of Arabidopsis grown on solid media supplemented with 0.5 mM GSH or CoM n = 3 biological replicates. Lack of asterisks indicates no p-value ≤ 0.05.

**Figure S2.** **Root growth of Arabidopsis is positively affected by CoM application** Root growth of Arabidopsis grown on solid media supplemented with 0.5 mM GSH of CoM n = 3 biological replicates. All asterisks indicate p-value ≤ 0.05.

**Figure S3.** **Intracellular Sulfur content of Arabidopsis is unaffected by CoM application** Measurements of internal S_34_ content of Arabidopsis grown on solid media supplemented with 0.5 mM GSH or CoM or 0.3 mM of MgSO_4_ x 7H_2_O n = 3 biological replicates. All asterisks indicate p-value ≤ 0.05.

**Figure S4.** **Growth of tobacco is increased by CoM application** 2^nd^ growth trial of tobacco grown to 21 days with spray application of CoM twice weekly with concentrations in mM listed. All asterisks indicate p-value ≤ 0.05.

**Figure S5.** **Stomatal conductance of tobacco is unaffected by CoM** **application** Stomatal conductance (*G_s_*) of tobacco grown to 21 days with spray application of 1mM CoM twice weekly as measured by LiCOR 6800 in response to increasing light. n = 6 biological replicates, lack of asterisks indicate no p-value ≤ 0.05.

**Table S1 Statistical analyses of Data**

| **Data** | **Test** | **p-value** |
| --- | --- | --- |
| Fig 1b Ctrl vs. 0.2 mM | Dunnett's multiple comparisons test | <0.0001 |
| Fig 1b Ctrl vs. 0.5 mM | Dunnett's multiple comparisons test | <0.0001 |
| Fig 1b Ctrl vs 1 mM | Dunnett's multiple comparisons test | <0.0001 |
| Fig 1b Ctrl vs 5 mM | Dunnett's multiple comparisons test | <0.0001 |
| Fig 1d Wet mass Ctrl vs GSH | Dunnett's multiple comparisons test | 0.0072 |
| Fig 1d Wet mass Ctrl vs CoM | Dunnett's multiple comparisons test | <0.0001 |
| Fig 1d Dry mass Ctrl vs GSH | Dunnett's multiple comparisons test | 0.0097 |
| Fig 1d Dry mass Ctrl vs CoM | Dunnett's multiple comparisons test | <0.0001 |
| Fig 1e -S vs -S+CoM | Dunnett's multiple comparisons test | <0.0001 |
| Fig 2a Ctrl vs 1 mM | Dunnett's multiple comparisons test | 0.0090 |
| Fig 2b Ctrl vs 1 mM | Dunnett's multiple comparisons test | 0.0060 |
| Fig 2b Ctrl vs 2.5 mM | Dunnett's multiple comparisons test | 0.0001 |
| Fig 2b Ctrl vs 5 mM | Dunnett's multiple comparisons test | 0.0003 |
| Fig 2c F_v_ Ctrl vs 1mM | Dunnett's multiple comparisons test | <0.0001 |
| Fig 2c F_v_ Ctrl vs 2.5 mM | Dunnett's multiple comparisons test | <0.0001 |
| Fig 2c F_v_ Ctrl vs 5 mM | Dunnett's multiple comparisons test | 0.0031 |
| Fig 2c F_m_ Ctrl vs 1mM | Dunnett's multiple comparisons test | <0.0001 |
| Fig 2c F_m_ Ctrl vs 2.5 mM | Dunnett's multiple comparisons test | <0.0001 |
| Fig 2c F_p_ Ctrl vs 1 mM | Dunnett's multiple comparisons test | <0.0001 |
| Fig 2c F_p_ Ctrl vs 2.5 mM | Dunnett's multiple comparisons test | <0.0001 |
| Fig 2d ɸPSII | 2-Way ANOVA (CoM concentration) | <0.0001 |
| Fig 2d ɸPSII | 2-Way ANOVA (Time) | <0.0001 |
| Fig 2d ɸPSII | 2-Way ANOVA (Interaction of concentration w/ Time) | <0.0001 |
| Fig 2d PSII_lightasym_ Ctrl vs 1 mM | Dunnett's multiple comparisons test | 0.0158 |
| Fig 2d PSII_lightasym_ Ctrl vs 2.5 mM | Dunnett's multiple comparisons test | 0.0014 |
| Fig 2d PSII_lightasym_ Ctrl vs 5 mM | Dunnett's multiple comparisons test | 0.0395 |
| Fig 2d PSII_end_ Ctrl vs 2.5 mM | Dunnett's multiple comparisons test | 0.0001 |
| Fig 2d PSII_end_ Ctrl vs 5 mM | Dunnett's multiple comparisons test | 0.0032 |
| Fig 2e ɸNPQ | 2-Way ANOVA (CoM concentration) | <0.0001 |
| Fig 2e ɸNPQ | 2-Way ANOVA (Time) | <0.0001 |
| Fig 2e ɸNPQ | 2-Way ANOVA (Interaction of concentration w/ time) | <0.0001 |
| Fig 2e NPQ_max_ Ctrl vs 1 mM | Dunnett's multiple comparisons test | 0.0058 |
| Fig 2e NPQ_max_ Ctrl vs 2.5 mM | Dunnett's multiple comparisons test | 0.0015 |
| Fig 2e NPQ_max_ Ctrl vs 5 mM | Dunnett's multiple comparisons test | 0.0004 |
| Fig 2e NPQ_rel_ Ctrl vs 1 mM | Dunnett's multiple comparisons test | 0.0360 |
| Fig 2e NPQ_rel_ Ctrl vs 2.5 mM | Dunnett's multiple comparisons test | <0.0001 |
| Fig 2e NPQ_rel_ Ctrl vs 5 mM | Dunnett's multiple comparisons test | <0.0001 |
| Fig 2e NPQ_end_ Ctrl vs 2.5 mM | Dunnett's multiple comparisons test | 0.0030 |
| Fig 2e NPQ_end_ Ctrl vs 5 mM | Dunnett's multiple comparisons test | 0.0221 |
| Fig 3a 28 days Ctrl vs 0.5 mM | Dunnett's multiple comparisons test | 0.0055 |
| Fig 3a 35 days Ctrl vs 0.5 mM | Dunnett's multiple comparisons test | 0.0037 |
| Fig 3b 0 mM vs 2 mM | Dunnett's multiple comparisons test | 0.0153 |
| Fig 3c 28 days 0 mM vs 3 mM | Dunnett's multiple comparisons test | 0.0060 |
| Fig 3c 35 days 0 mM vs 3 mM | Dunnett's multiple comparisons test | 0.0050 |
| Fig 3d Ctrl vs 3 mM | Dunnett's multiple comparisons test | 0.0025 |
| Fig 3e Ctrl 1mM | Dunnett's multiple comparisons test | <0.0001 |
| Fig S1 Ctrl vs GSH | Dunnett's multiple comparisons test | 0.0856 |
| Fig S1 Ctrl vs CoM | Dunnett's multiple comparisons test | >0.9999 |
| Fig S2 Ctrl vs CoM | Dunnett's multiple comparisons test | <0.0001 |
| Fig S3 Ctrl vs SO_4_ | Dunnett's multiple comparisons test | <0.0001 |
| Fig S4 Wet Mass Ctrl vs 1mM | Dunnett's multiple comparisons test | 0.0130 |
| Fig S5 G_s_ | 2-way ANOVA (Concentration) | 0.6817 |
| Fig S5 G_s_ | 2-way ANOVA (Light) | <0.0001 |
| Fig S5 G_s_ | 2-way ANOVA (Interaction of Concentration w/ Light) | 0.9686 |
